# CRISPR/Cas14a Combined with RPA for Visual Detection of Marek’s Disease Virus

**DOI:** 10.1101/2025.08.21.671677

**Authors:** Zhi-Jian Zhu, Meng-Li Cui, Yu Liu, Xi-Qiao Yao, Ming-Cheng Wang, Jun-He Liu, Jin-Feng Li, En-Zhong Li

## Abstract

Marek’s disease (MD), a highly contagious avian immunosuppressive disorder caused by the α-herpesvirus MDV-1, poses a significant threat to poultry health. The development of rapid visual detection methods capable of distinguishing epidemic MDV-1 strains from vaccine strains is crucial for early disease warning, vaccine efficacy evaluation, and precise disease control. We developed a novel isothermal detection system that integrates recombinase polymerase amplification (RPA) with CRISPR/Cas14a technology for the visual identification of epidemic MDV-1 strains. This method operates at a constant temperature of 37°C and allows for either real-time analysis or endpoint visual readout without the need for complex instrumentation. Our results showed no cross-reactivity with Newcastle disease virus (NDV), infectious bursal disease virus (IBDV), MDV-1 vaccine strains, or herpesvirus of turkeys (HVT). Plasmid DNA standards were used to determine the sensitivity of the assay and the detection limit was 24.6 copies/μL. Clinical evaluation using 24 field samples confirmed that the method successfully identified all MDV-positive cases, demonstrating its diagnostic reliability. In conclusion, we have developed a rapid, instrument-free, and highly specific nucleic acid detection platform for MDV-1 by combining the sensitivity of RPA with the specificity of CRISPR/Cas14a technology, offering promising potential for field-based diagnostics and disease surveillance.

**IMPORTANCE:** Marek’s disease virus (MDV-1) is a highly contagious and economically important avian pathogen. Existing diagnostic methods are unable to reliably distinguish between epidemic and vaccine strains in field settings, which hampers effective surveillance and evaluation of vaccination programs. To address this challenge, we developed a portable isothermal detection assay that combines recombinase polymerase amplification (RPA) with CRISPR/Cas14a technology. This approach enables highly sensitive (24.6 copies/μL) and specific visual detection of epidemic MDV-1 strains without cross-reactivity with vaccine strains or related viruses. The assay demonstrated 100% agreement with reference methods when validated using clinical samples. As a cost-effective and instrument-free method, it offers a practical solution for rapid on-site diagnosis, facilitating enhanced outbreak control and improved poultry health management globally.

---

Marek’s disease (MD) is a highly contagious and economically significant avian lymphoproliferative disorder caused by Marek’s disease virus (MDV)(1). It is characterized by immunosuppression, systemic organ dysfunction, and the development of lethal neoplastic transformations in infected chickens, resulting in substantial economic losses to the global poultry industry (2, 3). Based on taxonomic classification, MDV comprises three serotypes: *Gallid herpesvirus 2* (GaHV-2; MDV-1), *Gallid herpesvirus 3* (GaHV-3; MDV-2), and *Meleagrid herpesvirus 1* (MeHV-1; HVT)(4). Importantly, only field strains of MDV-1 possess oncogenic properties, capable of inducing malignant lymphomas in susceptible chicken populations. Currently, control measures for MD primarily rely on vaccination. The principal vaccine strains used for immunoprophylaxis include CVI988/Rispens and mMDV814, both derived from MDV-1, as well as FC-126, which is derived from HVT(5). Among these, CVI988/Rispens is recognized as the gold-standard attenuated vaccine due to its high efficacy against virulent (v MDV), very virulent (vv MDV), and very virulent plus (vv+ MDV) strains, thus being widely adopted in global MD vaccination programs. However, continuous immune pressure exerted by vaccines has contributed to the evolution of MDV virulence, leading to an increased frequency of vaccine breakthroughs(6, 7). MD outbreaks have become more prevalent in recent years, presenting significant challenges to effective disease control. Therefore, rapid and accurate differential diagnosis of suspected MD cases is essential to enable timely intervention by poultry producers and minimize economic losses.

Common diagnostic methods for Marek’s disease (MD) detection include virus isolation and culture, serological assays, and molecular testing. Virus isolation and identification involve complex and time-consuming procedures, with results often influenced by various confounding factors—particularly interference from vaccine strains. Serological diagnosis, on the other hand, is limited in its ability to reliably differentiate between field strains and vaccine-derived viruses in both infected individuals and clinical cases. Molecular techniques based on nucleic acid amplification, such as polymerase chain reaction (PCR), offer high-throughput and rapid detection capabilities, providing significantly shorter turnaround times and greater accuracy compared to traditional methods. These advantages position PCR as a crucial tool in MD diagnostics, effectively addressing the shortcomings of alternative approaches(8, 9). However, PCR-based detection requires costly equipment and highly trained personnel, which restricts its use in resource-limited settings that lack advanced laboratory infrastructure.

To meet the demands of field diagnostics outside laboratory settings, researchers have developed isothermal nucleic acid amplification techniques, such as loop-mediated isothermal amplification (LAMP) and recombinase polymerase amplification (RPA). RPA technology offers several advantages, including rapid amplification, high sensitivity, strong specificity, and the ability to operate at a constant temperature within the range of 37-42°C(10). Since its introduction, RPA has been widely applied in the detection of various viruses, such as severe acute respiratory syndrome coronavirus (SARS-CoV)(11), monkeypox virus (MPXV)(12), and african swine fever virus (ASFV)(13). Zeng et al. designed specific primers and probes targeting the *meq* gene of MDV for detection, achieving high specificity and sensitivity(14). However, RPA’s relatively high tolerance for primer binding site mismatches may result in nonspecific amplification among closely related species, limiting its ability to distinguish between vaccine strains such as CVI988/Rispens. Given this limitation, combining rapid RPA amplification with the precise gene-editing capabilities of CRISPR/Cas systems has emerged as a promising direction in the development of current and future pathogen diagnostic technologie(15–17).

The CRISPR/Cas system, as a revolutionary gene-editing tool, offers a variety of effector proteins that provide novel technological approaches for molecular detection(18). Among these, CRISPR/Cas14 represents the smallest RNA-guided nuclease identified to date within the CRISPR/Cas family, with the Cas14a effector protein typically exhibiting a relative molecular mass ranging from 40 to 70 kDa(19). Guided by sgRNA, Cas14a binds and cleaves single-stranded DNA (ssDNA) targets in a PAM-independent manner. Upon target recognition, Cas14a activates its trans-cleavage activity, enabling nonspecific degradation of ssDNA molecules. Due to its PAM-independent recognition mechanism, Cas14a requires high sequence fidelity—single-base mismatches can significantly reduce its cleavage efficiency. This characteristic endows Cas14a with exceptional sensitivity in detecting single-nucleotide polymorphisms (SNPs). In recent years, the integration of Cas14a’s trans-cleavage activity with diverse signal amplification strategies has enabled rapid and sensitive pathogen detection, highlighting its promising potential in diagnostic applications(20, 21). In this study, we developed an enhanced Cas14a-based detection method by combining RPA, λ exonuclease treatment, and the collateral activity of CRISPR/Cas14a, enabling sensitive, specific, equipment-free, and visually interpretable detection of MDV-1 through targeting of the conserved *meq* gene (Fig. 1).

**FIG 1.**
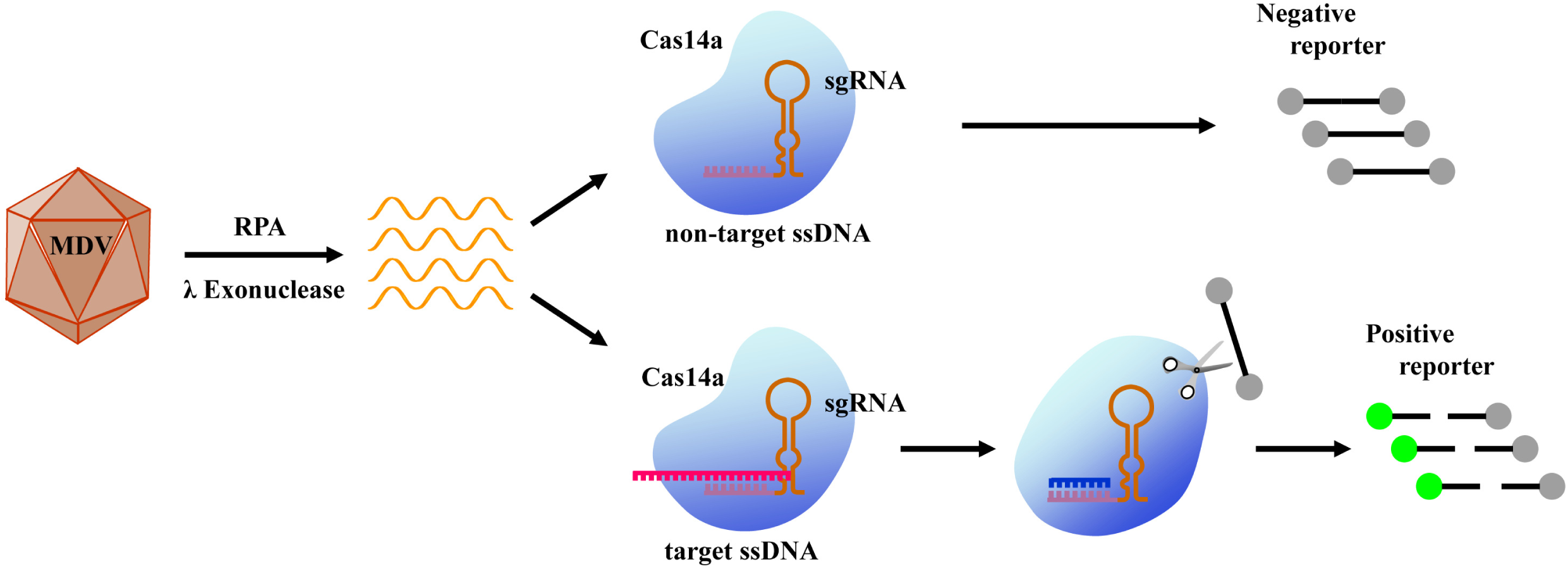
Schematic illustration of the MDV detection method utilizing the integration of RPA and CRISPR/Cas14a technologies.

## MATERIALS AND METHODS

### Viruses and clinical samples

The MDV-1 Md5 strain was obtained from the laboratory stock. The MDV-1 vaccine strain CVI988/Rispens and turkey herpesvirus (HVT) Fc-126 are commercially available vaccines provided by Merck Animal Health. The Newcastle disease virus (NDV) was supplied by Zhumadian Animal Disease Prevention and Quarantine Center, and the infectious bursal disease virus (IBDV) B87 was purchased from the Guangdong epidemiology prevention and control center. A total of 24 tissue samples were obtained from diverse poultry farms situated in Henan province. During the sample collection procedure, the animals were humanely euthanized via intraperitoneal injection of pentobarbital sodium at a dosage of 150 mg/kg. The collection of animal tissue samples was conducted in accordance with the guidelines approved by the Huanghuai University Experimental Animal Ethics Committee (NO. 20240306012).

### DNA/RNA extraction

The nucleic acid of viruses and clinical samples were extracted using the TIANamp Virus DNA/RNA kit (Tiangen Biotech, Beijing, China) according to the manufacturer’s instruction and stored at −80 °C until used.

### Generation of DNA standard

The open reading frame (ORF) of the Meq gene from the MDV-1 epidemic strain (international reference strain Md5, GenBank Acc. No.: AF243438.1) and the vaccine strain CVI988/Rispens (GenBank Acc. No.: DQ530348.1) was amplified using the forward primer 5’-TGCTGGAATGTTAAGAATAAATTCCGCAC-3’ and the reverse primer 5’-TTATCTCATACTTCGGAACTCCTGG-3’, respectively. The S-meq and L-meq genes amplified from Md5 and CVI988/Rispens were purified using the EasyPure® PCR Purification Kit (Tiangen Biotech, Beijing, China) and subsequently cloned into the pMD-19T vector (Takara, Dalian, China), resulting in the recombinant plasmids pMD-19T-S-meq and pMD-19T-L-meq. Positive clones were selected and sequenced for further processing. Plasmids were extracted using the SanPrep Plasmid Mini-Preparation Kit (Sangon Biotech, Shanghai, China), according to the manufacturer’s instructions. DNA concentration was measured by spectrophotometry, and the copy number was calculated prior to being used as a DNA standard for sensitivity analysis.

### RPA primer design and screening

Twenty-one nucleotide sequences of the reference MDV Meq genes were aligned using DNASTAR software to identify conserved regions (Fig. S1). Based on the highly conserved G→T single-nucleotide polymorphism at position 211 of the Meq gene between vaccine and epidemic strains, we designed RPA primer pairs specifically targeting this locus using Primer Premier 5.0 software, ultimately obtaining three primer pairs (comprising three forward and three reverse primers). All reverse primers had phosphorylated 5’ ends. The RPA primers were synthesized by Sangon Biotech (Table 1).

**TABLE 1.**
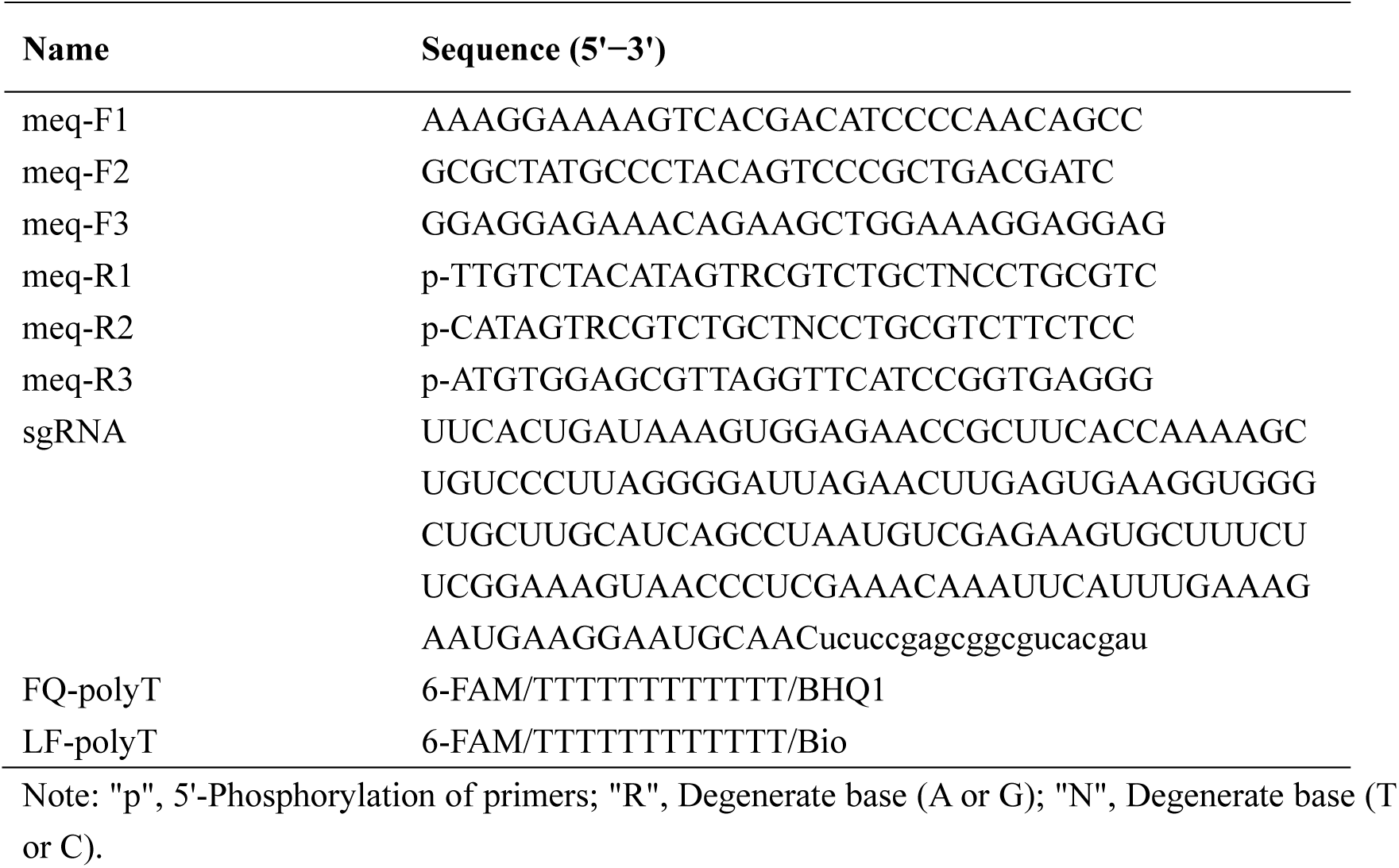
Sequence of RPA primers, sgRNA FQ-polyT and LF-polyT in this study

The RPA reaction was conducted using a TwistAmp® Basic kit (TwistDX, Cambridge, United Kingdom). A 1 μl aliquot of positive control plasmid pMD19-T-S-meq served as as the template and was amplified with different primer combinations (Table 1), according to the manufacturer’s instructions. The reaction was carried out in a 50 μl system at 37°C for 20 minutes. The resulting RPA amplicons were separated by electrophoresis on a 1% agarose gel for analysis.

### sgRNA preparation and Cas14a detection

For sgRNA preparation targeting the single-nucleotide polymorphism at position 211 of the Meq gene, DNA templates of sgRNA were appended with the T7 promoter sequence and synthesized as primers by Sangon Biotech (Table 1). The two primers were annealed into double-stranded DNA using the Annealing Buffer for DNA Oligos (Beyotime). The double-stranded DNA was purified through gel extraction. Using the T7 high efficiency transcription kit (TransGen Biotech), the DNA was transcribed into sgRNA via overnight incubation at 37°C. The resulting sgRNA was then purified using the EasyPure® RNA purification kit (TransGen Biotech) following the manufacturer’s protocol and stored at −80°C.

Cas14a detection was carried out according to the manufacturer’s instructions (Sangon Biotech, Shanghai, China). The 50 μl Cas14a reaction mixture contained 5 μl RPA amplification products, 25 nM sgRNA, 50 nM Cas14a, 2 μl RNase inhibitor (New England Biolabs), 100 nM ssDNA reporter (Table 1), 0.6 μl λ Exonuclease (Lambda Exonuclease), and Cas14a detection buffer (1×). Fluorescence kinetics were monitored at 37°C over a 2-hour period using an excitation wavelength of 485 nm and an emission wavelength of 520 nm, with fluorescence readings recorded every 4 minutes.

### Cas14a detection with lateral Flow Assay

For lateral flow-based Cas14a detection, the ssDNA reporter was replaced with a synthesized FAM-ssDNA-biotin reporter at a final concentration of 100 nM in a 50 μL Cas14a reaction system. The Cas14a reaction was performed at 37°C for 1 hour. Subsequently, the CRISPR Cas12/13 HybriDetect Dipstick (WARBIO) was immersed into the solution. Following a 5-minute incubation period, images of the dipsticks were captured using a camera.

### Clinical sample detection

The performance of the RPA-CRISPR/Cas14a assay was assessed using a panel of 24 clinical samples. For comparative analysis, all samples were simultaneously analyzed using a PCR assay targeting the Meq gene. The PCR reaction was carried out in a 20 μL volume consisting of 10 μL of 2 × PCR Mix (Takara, Dalian, China), 2 μM each of the forward primer (5′-TGCTGGAATGTTAAGAATAAATTCCGCAC-3′) and reverse primer (5′-TTATCTCATACTTCGGAACTCCTGG-3′), and 25 ng of DNA template. Amplification products were analyzed by electrophoresis on a 1% agarose gel.

## RESULTS

### Optimal RPA primer set selection

In this study, three forward and three reverse primers were designed and systematically evaluated across all possible combinations, yielding nine distinct primer sets. These were assessed using real-time RPA. The results showed that all primer combinations effectively recognized and amplified the target sequence of MDV. Comparative analysis demonstrated that the meq-F1/R3 primer pair demonstrated significantly higher amplification efficiency and specificity compared to the other combinations (Fig. 2). Consequently, this primer set was identified as the optimal RPA primer system and was utilized in subsequent experiments.

**FIG 2.**
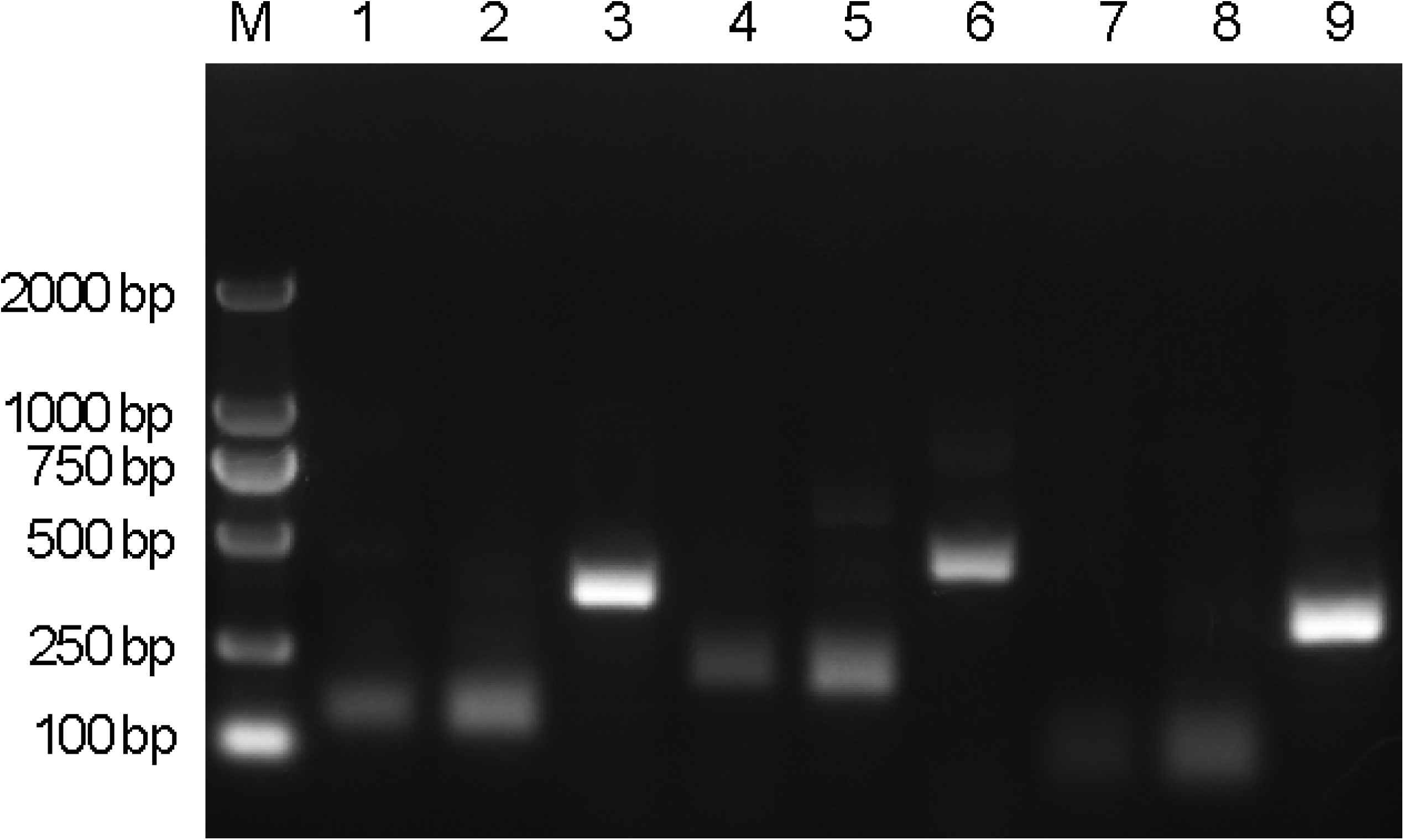
Agarose gel electrophoresis analysis of RPA-amplified products targeting the Meq gene of MDV-1 strain. M: DNA Marker; 1: meq-F1/meq-R1; 2: meq-F1/meq-R2; 3: meq-F1/meq-R3; 4: meq-F2/meq-R1; 5: meq-F2/meq-R2; 6: meq-F2/meq-R3; 7: meq-F3/meq-R1; 8: meq-F3/meq-R2; 9: meq-F3/meq-R3.

### Validation of the sgRNA-Cas14a fluorescence detection

To evaluate the sensitivity and specificity of the designed sgRNA in recognizing the Meq gene target sequence among prevalent MDV strains, the complete Cas14a reaction system and the Cas14a reaction system lacking sgRNA were comparatively analyzed. The positive control plasmid pMD-19T-S-meq was used as the template in the RPA assay, and its amplification products were introduced into either the complete Cas14a reaction system or the system without sgRNA for fluorescence kinetic analysis. As shown in figure 3, the relative fluorescence units (RFU) in the complete Cas14a system increased rapidly over time until the peak value, whereas no significant fluorescence signal was detected in the system without sgRNA. To further verify the specificity of sgRNA-mediated recognition of the Meq target sequence, RPA products derived from the negative control plasmid pMD-19T-L-meq were subjected to the same fluorescence-based Cas14a detection system. Notably, only the pMD-19T-S-meq plasmid exhibited a time-dependent increase in RFU, while both the negative control and pMD-19T-L-meq groups remained non-reactive (Fig. 3). These results demonstrate that the sgRNA-Cas14a fluorescence detection system enables highly specific identification of MDV-1 epidemic strains.

**FIG 3.**
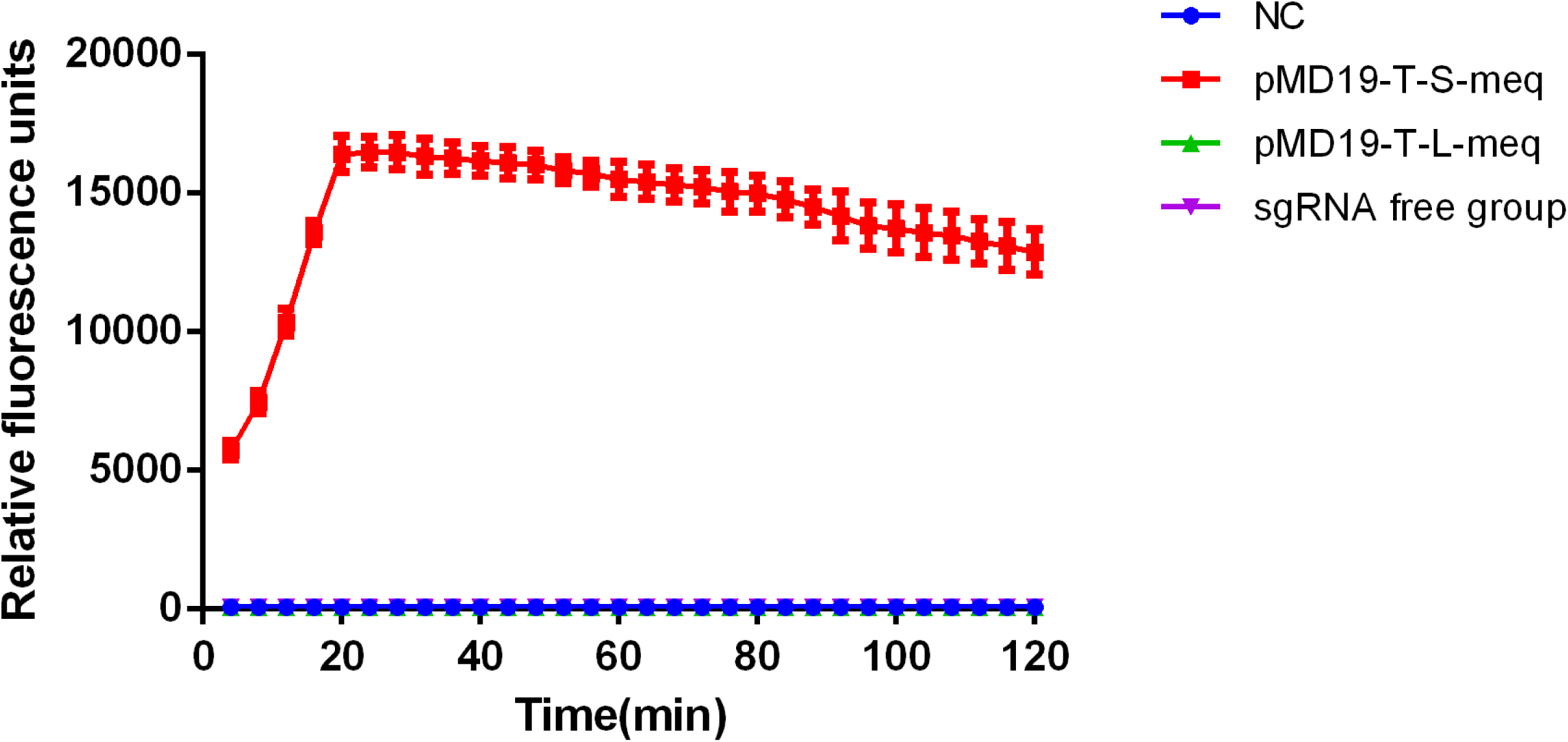
Analysis of positive standard plasmid pMD-19T-S-meq by RPA-CRISPR/Cas14a fluorescence detection. (n = 3 technical replicates; values represent mean ± SEM).

### Specificity and sensitivity of the enhanced Cas14a fluorescence detection

The specificity of the enhanced Cas14a fluorescence detection was assessed using the MDV international standard virulent strain (Md5), vaccine strains (CVI988/Rispens and HVT), as well as other avian viruses including Newcastle disease virus (NDV) and infectious bursal disease virus (IBDV). As shown in figure 4A, a rapid and strong fluorescent signal was observed exclusively for the Md5 strain. In contrast, no detectable fluorescence was generated in response to the MDV-1 vaccine strains, other tested viruses, or the negative controls. These findings indicate that the proposed detection method possesses high specificity and does not exhibit cross-reactivity with other pathogens, thereby enabling accurate differentiation of MDV-1 epidemic strains.

**FIG 4.**
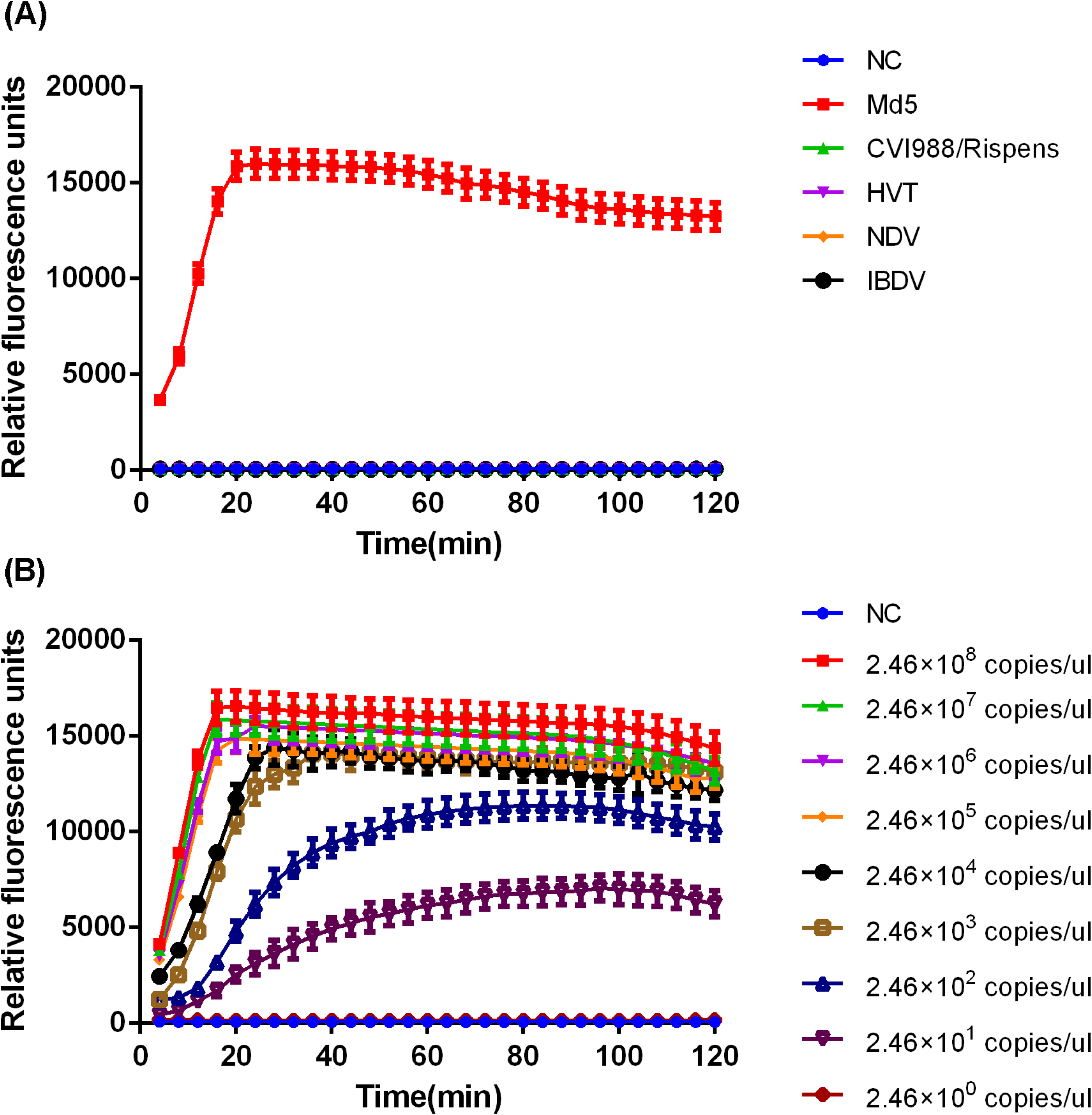
Specificity and sensitivity of RPA-CRISPR/Cas14a fluorescence detection. (A) Specificity of RPA-CRISPR/Cas14a fluorescence detection. (n = 3 technical replicates; values represent mean ± SEM). (B) Sensitivity of RPA-CRISPR/Cas14a fluorescence detection. (n = 3 technical replicates; values represent mean ± SEM).

The sensitivity of the enhanced Cas14a fluorescence detection assay was evaluated using a 10-fold serial dilution of the pMD-19T-S-meq positive standard plasmid. As illustrated in figure 4B, the system was capable of detecting target DNA across seven orders of magnitude, ranging from 2.46 × 10⁸ copies/μL to 2.46 × 10^1^ copies/μL. These data showed that the limit of detection for the enhanced Cas14a fluorescence detection method is 24.6 copies/μl.

### Validation of the enhanced Cas14a lateral flow detection

To enable rapid on-site detection with visual readout, we developed an enhanced lateral flow detection (LFD) system based on the Cas14a protein. This assay employs a FAM-labeled ssDNA-biotin reporter for Cas14a-mediated target recognition. In negative samples, gold-conjugated anti-FAM antibodies bind to intact reporter molecules, forming complexes that are captured by biotin ligands at the control line. In positive samples, Cas14a-mediated cleavage of the reporter redirects the gold antibody-FAM complexes to the test line, accompanied by reduced control line signal intensity. When applied to detect pMD-19T-S-meq and pMD-19T-L-meq plasmids, the assay specifically identified pMD-19T-S-meq, as indicated by a clearly visible test line (Fig. 5). These results confirm that the enhanced Cas14a-LFD enables visual discrimination of MDV-1 epidemic strains.

**FIG 5.**
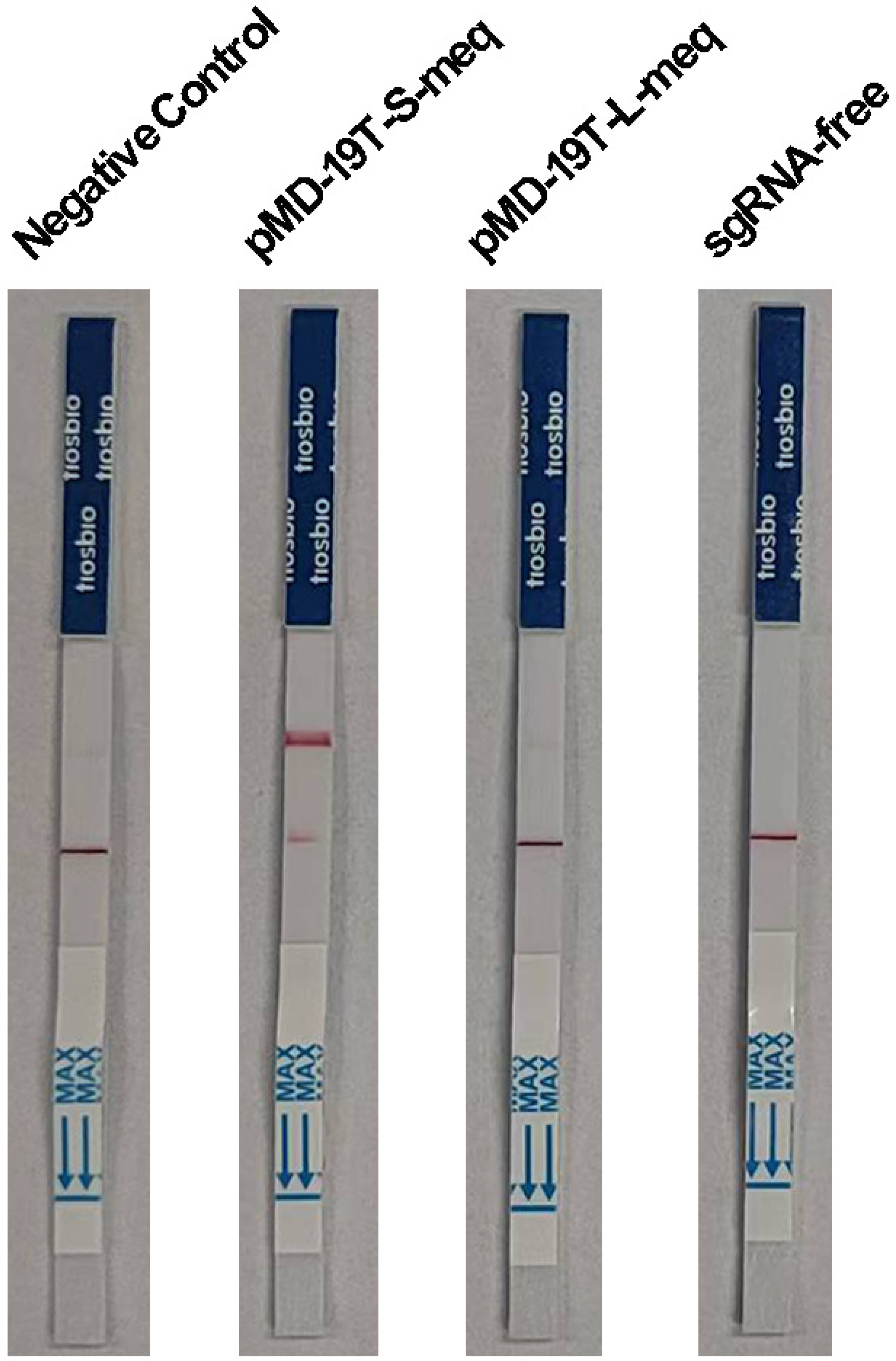
Analysis of positive standard plasmid pMD-19T-S-meq by RPA-CRISPR/Cas14a lateral flow detection.

### Specificity and sensitivity of the enhanced Cas14a lateral flow detection

To evaluate the specificity of the enhanced Cas14a-based lateral flow assay, various viruses, including the Md5 strain, vaccine strain CVI988/Rispens, HVT, NDV, and IBDV, were analysed. The results showed that positive signals were exclusively detected on the lateral flow strips corresponding to the Md5 strain (Fig. 6A). To assess the sensitivity of the enhanced Cas14a lateral flow detection method, 10-fold serial dilutions of the pMD-19T-S-meq plasmid described previously were tested (Fig. 6B). The detection limit was determined to be 24.6 copies/μL, which is consistent with the sensitivity observed in the enhanced Cas14a fluorescence detection method. Therefore, the enhanced Cas14a lateral flow assay can be effectively utilized for specific and sensitive detection of MDV-1 epidemic strains.

**FIG 6.**
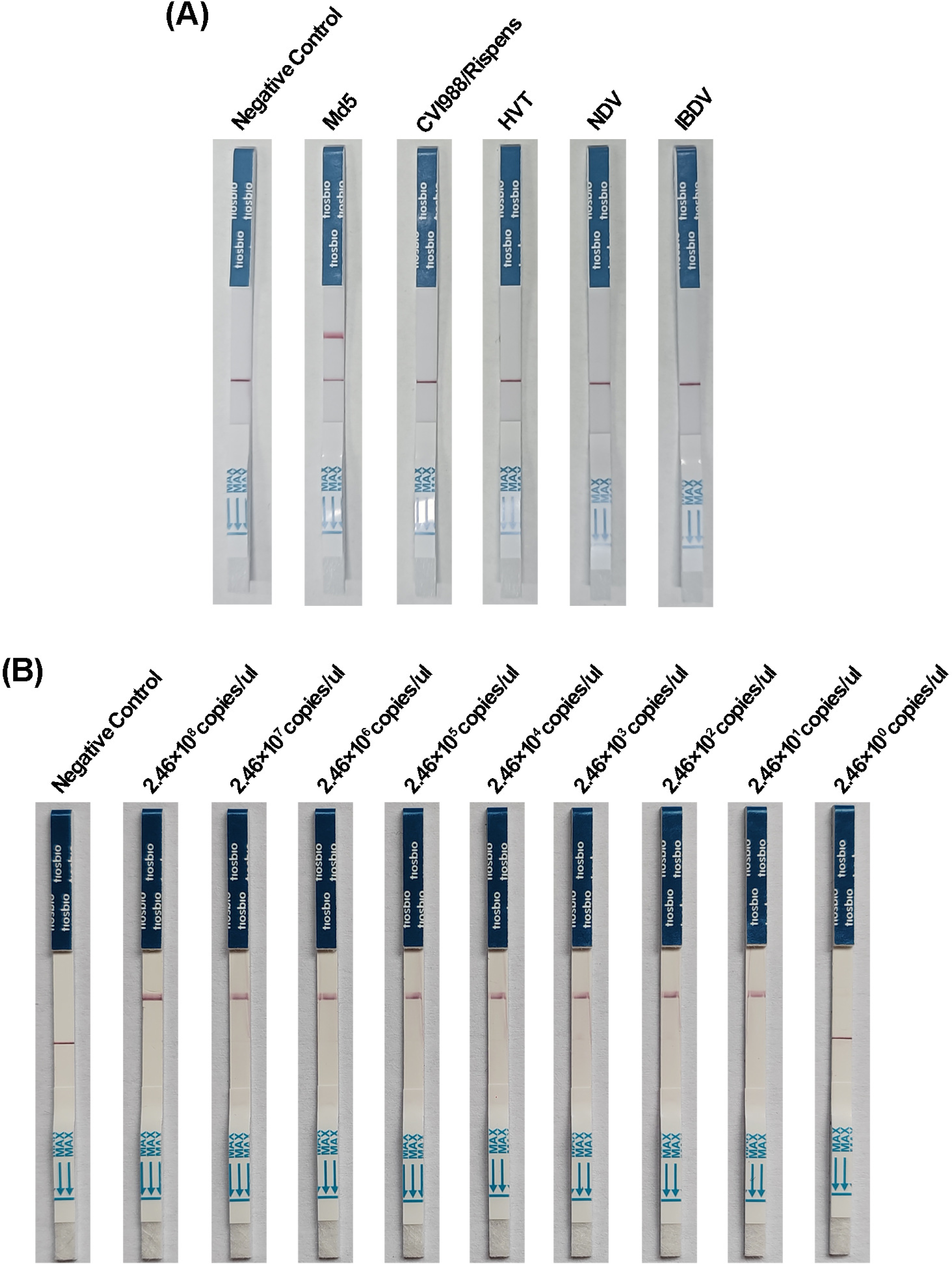
Specificity and sensitivity of RPA-CRISPR/Cas14a lateral flow detection. (A) Specificity of RPA-CRISPR/Cas14a lateral flow detection. (B) Sensitivity of RPA-CRISPR/Cas14a lateral flow detection

### Enhanced Cas14a detection in clinical samples

To evaluate the clinical applicability of the enhanced Cas14a detection system, 24 tissue samples were collected from various farms and simultaneously tested using three methods: the enhanced Cas14a fluorescence assay, the enhanced Cas14a lateral flow assay, and conventional PCR. Both Cas14a-based methods (fluorescence and lateral flow) identified 17 samples as MDV-1-positive and 7 as MDV-1-negative. For validation purposes, PCR was performed using Meq gene-specific primers. The PCR results were fully consistent with those obtained from both enhanced Cas14a assays (Table 2). These findings indicate that the enhanced Cas14a fluorescence and lateral flow assays exhibit strong potential as on-site diagnostic tools for detecting MDV-1 epidemic strains in clinical settings.

**TABLE 2.** Detection in clinical samples by RPA-CRISPR/Cas14a detection and PCR

| Assay | Number of samples |  |  |
| --- | --- | --- | --- |
|  | Positive | Negative | CVI988/814 |
| RPA-CRISPR/Cas14a fluorescence detection | 17 | 7 | / |
| RPA-CRISPR/Cas14a lateral flow detection | 17 | 7 | / |
| PCR | 17 | 7 | 10 |

## DISCUSSION

Nucleic acid-based detection techniques have emerged as powerful tools for MDV identification due to their high sensitivity. Among these, RPA has attracted increasing attention in MDV clinical diagnostics because of its minimal dependence on complex instrumentation and strict experimental conditions. However, RPA’s high tolerance to to base-pair mismatches limits its ability to distinguish pathogenic MDV strains from vaccine strains, such as CVI988 or mMDV814. The mandatory administration of MDV vaccines in poultry farming across China presents a significant challenge for the broad implementation of RPA-based MDV detection methods in clinical diagnostics. Thus, there is an urgent need to develop a novel MDV detection method that combines the high sensitivity of nucleic acid testing with improved specificity.

In this study, we developed a novel MDV nucleic acid detection system by integrating RPA and CRISPR/Cas14a technologies. Given that the Meq gene is a unique molecular marker of MDV-1 and its encoded protein plays a crucial role in viral oncogenicity and pathogenesis, we selected it as the detection target(22, 23).

During method development, we conducted a multiple sequence alignment of the Meq gene from 21 MDV reference strains in the NCBI database to identify highly conserved regions for RPA primer design. Notably, a G→T single nucleotide polymorphism (SNP) distinguishing vaccine strains from epidemic strains was identified within the conserved region, enabling the design of a strain-specific sgRNA. Performance evaluation demonstrated that the RPA-CRISPR/Cas14a system exhibited excellent detection characteristics for MDV-1 epidemic strains, with a sensitivity of 24.6 copies/μL. Compared with the RPA method reported by Zeng et al. [14], which has a detection limit of 100 copies per reaction, the proposed system showed comparable sensitivity while preserving the high sequence-specific recognition capability inherent to CRISPR technology.

The RPA-CRISPR/Cas14a detection system exhibited strong applicability in clinical sample validation. Analysis of 24 clinical specimens collected from poultry farms across Henan province in China, revealed that 17 tissue samples were positive for MDV-1, showing complete concordance with conventional PCR results and thereby confirming the reliability of the system for the clinical diagnosis of Marek’s disease. Importantly, the Cas14a protein not only enables fluorescence-based detection but can also be adapted for visual readout using lateral flow dipsticks, offering dual advantages of laboratory-grade quantification and field-deployable rapid screening. Furthermore, the dual specificity conferred by RPA amplification coupled with sgRNA-mediated targeting ensures reliable discrimination between vaccine strains and wild-type viruses. This crucial feature significantly enhances the system’s value in avian disease surveillance, addressing a critical need in the poultry industry for rapid and differential diagnosis of pathogenic MDV strains.

In conclusion, a novel visual nucleic acid detection method based on RPA-CRISPR/Cas14a technology was established for MDV. This newly developed assay provides a reliable alternative for MDV detection, requiring minimal equipment and showing great potential for application in resource-limited settings.

## ACKNOWLEDGMENTS

We are sincerely grateful to the Institute for Animal Health, Henan Academy of Agricultural Sciences for their valuable cooperation and support throughout the research process.

This work was supported by the Key Scientific Research Projects of Universities in Henan (no. 22A310017).

## FUNDING

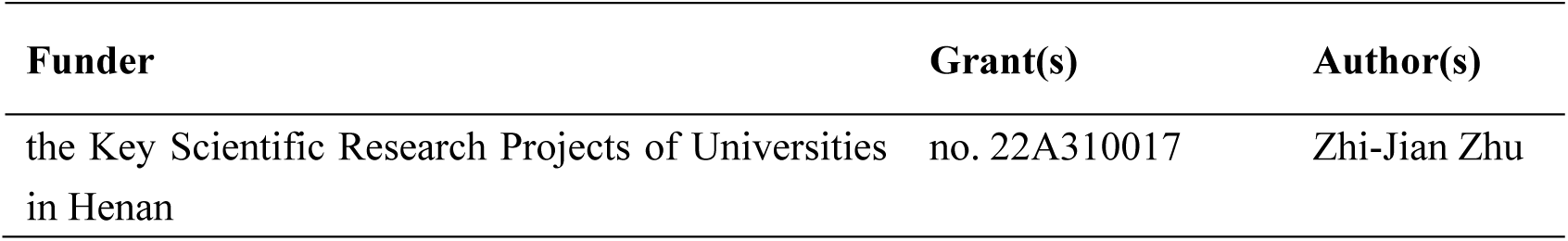

### AUTHOR CONTRIBUTIONS

Z.Z.J. and L.E.Z. contributed to the design of this work. Z.Z.J., L.E.Z and L.J.F. contributed to the data analysis and interpretation. Z.Z.J., C.M.L., L.Y., Y.X.Q., W.M.C. and L.J.H. contributed to the result validation. Z.Z.J., L.E.Z, C.M.L., L.Y., Y.X.Q. and L.J.F contributed to drafting and editing of this article. All authors have read and agreed to the published version of the manuscript.

### DATA AVAILABILITY

The data generated or analyzed during this study are available from the corresponding author upon reasonable request.

### ETHICS APPROVAL

All experimental protocols were approved prior to the study by the Huanghuai University Experimental Animal Ethics Committee (NO. 20240306012).

### ADDITIONAL FILES

The following material is available online.

